# Genomic prediction accounting for genotype by environment interaction offers an effective framework for breeding simultaneously for adaptation to an abiotic stress and performance under normal cropping conditions in rice

**DOI:** 10.1101/257808

**Authors:** M Ben Hassen, J Bartholomé, G Valè, TV Cao, N Ahmadi

**Affiliations:** CIRAD, UMR AGAP, F-34398 Montpellier, France; AGAP, Univ Montpellier, CIRAD, INRA, Montpellier SupAgro, Montpellier, France; CREA-Council for Agricultural Research and Economics, Research Center for Cereal and Industrial Crops, S. S. 11 to Torino Km 2.5, Vercelli, 13100, Italy

**Keywords:** Rice, genomic selection, progeny prediction, G×E interaction, alternate wetting and drying (AWD)

## Abstract

Developing rice varieties adapted to alternate wetting and drying water management is crucial for the sustainability of irrigated rice cropping systems. Here we report the first study exploring the feasibility of breeding rice for adaptation to alternate wetting and drying using genomic prediction methods that account for genotype by environment interactions. Two breeding populations (a reference panel of 284 accessions and a progeny population of 97 advanced lines) were evaluated under alternate wetting and drying and continuous flooding management systems. The accuracy of genomic prediction for response variables (index of relative performance and the slope of the joint regression) and for multi-environment genomic prediction models were compared. For the three traits considered (days to flowering, panicle weight and nitrogen-balance index), significant genotype by environment interactions were observed in both populations. In cross validation, prediction accuracy for the index was on average lower (0.31) than that of the slope of the joint regression (0.64) whatever the trait considered. Similar results were found for across population validation (progeny validation). Both cross-validation and progeny validation experiments showed that the performance of multi-environment models predicting unobserved phenotypes of untested entrees was similar to the performance of single environment models with differences in accuracy ranging from - 6% to 4% depending on the trait and on the statistical model concerned. The accuracy of multi-environment models predicting unobserved phenotypes of entrees evaluated under both water management systems outperformed single environment models by an average of 30%. Practical implications for breeding rice for adaptation to AWD are discussed.

## Introduction

Rice is the world’s most important staple food and will continue to be so in the coming decades. In the future, the necessary increases in rice production to meet demand will have to come mainly from an increase in yield per unit of land, water and other resources (CGIAR Research Program on Rice 2016). At the same time, 15–20 million ha of rice lands will suffer some degree of water scarcity (Tuong and Bouman 2003; Mekonnen and Hoekstra 2016). The predicted increase in water scarcity threatens the sustainability of rice production (Rijsberman 2006). It is thus crucial to develop agronomic practices that reduce water use while maintaining or increasing yields. A concomitant challenge is to adapt rice varieties to these water-saving agronomic practices by improving their performance under water-limited conditions.

In recent decades, different water management systems have been developed with the aim of reducing water consumption by irrigated rice (Tuong et al. 2005; Yang et al. 2007). Among them, the alternate wetting and drying (AWD) system, in which paddy fields are subjected to intermittent flooding with dry periods managed by soil water potential measurements, is one of the most widely used (Linquist et al. 2015; Lampayan et al. 2015). A meta-analysis of 56 studies comparing AWD with continuous flooding (CF) reported an overall decrease in yield of about 5% (Carrijo et al. 2017). However, marked variations were observed mainly depending on the severity of the drying phase (i.e. the soil moisture at the end of each drying cycle) and on soil characteristics (Lampayan et al. 2015; Carrijo et al. 2017). Significant differences in genotypic responses to AWD, measured by changes in grain yield, have also been reported and attributed to modified biomass partitioning (Bueno et al. 2010). Root architectural traits such as the number of nodal roots and root dry weight at a depth of 10-20 cm 22-30 days after transplanting also significantly contribute to yield stability under AWD (Sandhu et al. 2017). Genome wide association analysis using a diversity panel revealed AWD-specific associations for several agronomic traits including days to flowering, plant height, tillering, and panicle and seed characteristics (Volante et al. 2017). Thus, rice adaptation to AWD appears to involve typical complex traits, whose improvement requires genome-wide breeding approaches that account for genotype by environment (G×E) interactions, i.e. the amplitude of the response of the genotypes to a shift from CF management to the AWD system.

In plant breeding, G×E interactions are usually assessed in multi-environment trials and expressed as a change in the relative performance of genotypes in different environments, with or without change in the ranking of the genotypes (Freeman 1973). G×E analysis plays a fundamental role in assessing genotype stability, in predicting the performance of untested genotypes and in maximizing response to selection. Statistical methods for assessing G×E interactions and estimating their sizes and opportunities to exploit them are widely discussed in the literature (Freeman 1973; Cooper et al. 1993;; Malosetti et al. 2013; Elias et al. 2016; de Leon et al. 2016). One of the earliest and most widely used methods is linear regression of the performance (often of yield) of the individual genotype on the mean performances of all genotypes evaluated in each test environment (Yates and Cochran 1938). The method, known as *joint regression analysis*, was further formalized by Eberhart and Russel (1966) to enable testing of the significance of deviation of individual regression from the general linear component of G×E. Most evaluations of the effect of the environment on performance undertaken for the purpose of plant breeding rely on multi-environmental field testing that represents target production environments or a target population of environments (Cooper and Hammer 1996). One specific case of G×E experiments is managed-environment trials that aim to <u>assess</u> the effect of particular genvironmental variables (e.g., abiotic stresses) or cropping practices (e.g. fertilizer, irrigation, etc.) that influence crop performance in the production environment concerned (Cooper and Hammer 1996). A still more specific case of G×E experiments is managed abiotic stress trials that aim to provide a measure of genotypic response to stress based on yield loss under stress compared with under normal conditions. Several indexes have been proposed to evaluate the stress intensity and genotypic response in such experiments, mainly in the context of selection for drought tolerance (Fischer and Maurer 1978; Rosielle and Hamblin 1981; Fischer et al. 2003).

With the advent of molecular markers, new G×E analysis methods have been developed based on linear mixed models that connect the differential sensitivity of genotypes to environments to particular regions of the plant genome and to specific biological mechanisms (Malosetti et al. 2004; Boer et al. 2007; van Eeuwijk et al. 2010). More recently, the potential of genomic selection (GS) to accelerate the pace of genetic gains in major field crops has encouraged the development of multi-environment models for genomic prediction. The first statistical framework using a linear mixed model to model G×E for the purpose of genomic prediction was proposed by Burgueño et al. (2012). It extended the single-trait, single-environment genomic best linear unbiased prediction (GBLUP) model to a multi-environment context. Jarquín et al. (2014) proposed a method of modeling interactions between a high-dimensional set of markers and environmental that incorporates genetic and environmental gradients, as random linear functions (reaction norm) of markers and environmental covariates, respectively. Lopez-Cruz et al. (2015) proposed a marker × environment interaction (M×E) GS model that can be implemented using regression of phenotypes on markers or using co-variance structures (a GBLUP-type model). Cuevas et al. (2016) further developed this approach by using a non-linear (Gaussian) kernel to model the G×E: the reproducing kernel Hilbert space with kernel averaging and the Gaussian kernel with the bandwidth estimated using an empirical Bayesian method. Crossa et al. (2016) extended the M×E model using priors that produce shrinkage (Bayesian ridge regression) or variable selection (BayesB), and reported better prediction performances for these models compared to single environment and across-environment models. The latest multi-environment genomic prediction models fall back on a Bayesian approach (Cuevas et al. 2017). Application of these methods to one maize and four wheat CIMMYT data sets showed that models with G×E always have higher prediction ability than single-environment models, regardless of the genetic correlation between environments. The predictive ability of these Bayesian methods was also generally better than that obtained with the G×E models proposed by Lopez-Cruz et al. (2015) and Cuevas et al. (2016), when applied to the same datasets.

In the present study, we evaluated the effect of AWD on the performance of two rice breeding populations: a reference panel and a population of advanced lines both genotyped with 32 k SNP markers. Our general objective was to explore the feasibility of genomic selection for the adaptation of rice to AWD in the framework of a pedigree breeding scheme. Our specific objectives were to: (i) access expression of the response of the above-mentioned populations to AWD compared to the CF irrigation system, and (ii) compare the performance of different genomic prediction models that include G×E interactions in answering the two well-known issues relevant in breeding programs: predicting unobserved phenotypes of untested lines and predicting unobserved phenotypes of lines that have been evaluated in some environments but not others. The two issues are analyzed in the context of intra-population prediction (cross-validation experiments), and across-populations prediction (progeny-validation), as the population of advanced lines was derived from biparental crosses between some of the members of the diversity panel.

## Material and method

### Field trial and phenotyping

The plant material used in this study comprised a reference population (RP) of 284 accessions belonging to the rice *japonica* subspecies, and a progeny population (PP) of 97 advanced (F_5_-F_7_) inbred lines. The RP is representative of the working collection of the Research Centre for Cereal and Industrial Crops (CREA), Vercelli, Italy. The PP was derived from bi-parental crosses involving 31 accessions of RP, using a pedigree breeding scheme. More information on the two populations is provided in Ben Hassen et al. (2017). The two populations were phenotyped separately for two consecutive years at the experimental station of the CREA (45°19’24.00”N; 8°22’26.28”E; 134 m asl.): in 2012 and 2013 for RP and in 2014 and 2015 for PP. In each year, the phenotyping experiment included two independent trials corresponding to the two water management systems tested: CF and AWD. For the conventional CF water management system, rice was dry seeded and the field was flooded with 10-15 cm of water at the 3-4 leaf stage (typically 30 days after sowing) and maintained flooded until mid-maturity. For the AWD, after initial flooding at the 3-4 leaf stage, the field was subjected to intermittent drying periods. The soil water potential was maintained above −30 kPa by gravity irrigation whenever the soil moisture reached this threshold. The soil water potential was monitored by a set of six tensiometers distributed throughout the field and inserted to a depth of 20 cm. For each population and each year, the two water management systems were conducted in two fields separated by a distance of about 100 m to avoid interference with respect to the water regime. The other soil characteristics were identical (sand 47.8%, loam 42.8%, clay 9.4%; pH-H2O 6.4). The experimental design, which was identical in the two conditions, was a complete randomized design with three replicates for RP and a complete randomized block design with three replicates for the PP. The target traits for both RP and PP were days to flowering (FL), panicle weight (PW), and the nitrogen balance index (NI) as described in Ben Hassen et al. (2017).

### Modeling of phenotypic data

Phenotypic data for each condition in the RP and the PP were analyzed using mixed models. In order to identify possible outliers among individual data points, a diagnostic analysis based on restricted likelihood distance was implemented, for details see Ben Hassen et al. (2017). This analysis led to the elimination of one accession in the RP in AWD 2012 and 2013 experiments, one data point for FL in AWD-2012, and one data point for PW in AWD-2013. The discarded data were considered as missing in the following steps of the analysis. The following mixed models were applied to obtain adjusted means per genotype:

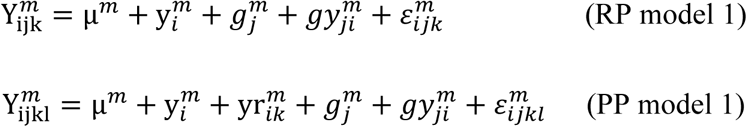

where Y^*m*^ is the observed phenotype for the water management system *m*; *μ*^*m*^ the overall mean; y^*m*^ the year as fixed effect; yr^*m*^ the within year replication as fixed effect; *g*^*m*^ the genotype as random effect, *gy*^*m*^ the interaction between genotype and year as random effect; and *ε*^*m*^ the residual. The analysis was performed with the *proc mixed* procedure of SAS 9.2 (SAS Institute, Cary NC, USA); the method of estimation for the variance components was the restricted maximum likelihood (REML). The formula by Holland et al. (2003) was used to estimate broad sense heritability (*H*^*2*^) as well as the corresponding standard error (SE) for each trait and each water management system in each population:

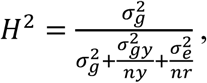

where 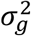, 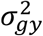 and 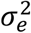 are the variance components associated with the genotype, the interaction between genotype and year and the residual, respectively. *ny* is the harmonic mean of the number of years per accession and *nr*, the harmonic mean of the number of plots across years per accession. Conditional coefficients of determination (R^2^) were also computed using the methodology described by Nakagawa and Schielzeth (2013). The adjusted means per water management system 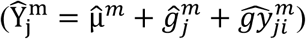 extracted from the model were used as phenotypes in the following steps.

For each trait, genetic correlations (*r_G_*) between values measured under the two water management systems were calculated (Cooper and DeLacy 1994; Cooper and Hammer 1996). The confidence interval of *r_G_* was obtained by using Fisher transformation of the estimated correlation 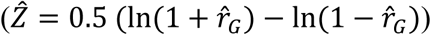, estimating the lower and upper bounds of 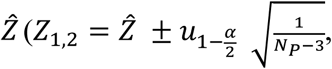, with *α =* 0.05, and *N_P_* = 284 and 97, for RP and PP respectively), and back transforming the 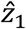 and 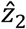 bounds into 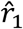 and 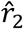. The ratio of correlated response to selection under continued flooding (CR_*CF*_) and the direct response under alternate watering and drying (DR_*AWD*_) was calculated as: 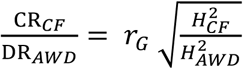 (Falconer 1989) where *r_G_* is the genotypic correlation defined above, and 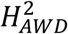 and 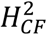 represent the heritability of the trait under AWD and CF, respectively.

In addition to models for each condition, a model gathering data from the two water management systems was also adjusted in order to test the significance of the interaction between water management and genotypes:

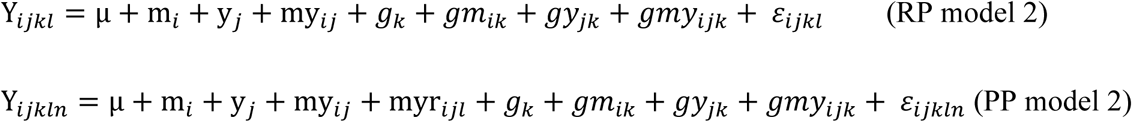

The same notation was used as for the model for each condition with additional fixed and random effects: *m* the water management as fixed effect; my the water management and year interaction as fixed effect; myr the replication within water management and year as fixed effect; *gm* the interaction between genotype and water management as random effect; and *gmy* the interaction between genotype, water management and year as random effect. The analyses were performed with the *proc mixed* procedure of SAS 9.2 (SAS Institute, Cary NC, USA) with REML.

### Evaluation of genotypic response to water management systems

The genotypic response to AWD water management was estimated in two ways using adjusted means. First, an index of relative performance was computed as follows: 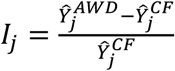, where 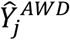 and 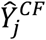 correspond to the adjusted means of accession *j* under AWD and CF water managements, respectively. This index was also calculated at population level to assess the intensity of stress caused by AWD water management compared to CF: *I* = 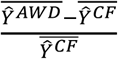 were 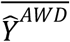 and 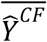 are the average performances of all genotypes within each population under AWD and CF, respectively. Second, the slope β_j_ was computed as defined in the joint regression equation: 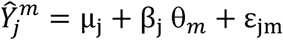, where 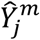 is the adjusted mean of the genotype *j* in the water management *m*; θ_m_ is the environmental index calculated as the mean value of all genotypes in water management *m*; *μ*_j_ is the intercept of the regression line of genotype j; and *ε*_*jm*_ is the residual.

### Genotypic data

The method used to genotype both RP and PP populations is detailed in Ben Hassen et al. (2017). It relies on the genotyping by sequencing protocol developed by Elshire et al. (2011). Sequencing was performed with a Genome Analyzer II (Illumina, Inc., San Diego, USA). The different steps of analysis (raw data filtering, sequence alignment, SNP calling and imputation) were performed with TASSEL v3.0 and the associated GBS pipeline (Glaubitz et al. 2014). A working set of 32,066 SNPs was obtained with a heterozygosity rate < 5% and minor allele frequency (MAF) > 5%.

### Statistical models for genomic prediction

#### Single environment models

To predict the genomic estimated breeding values within each water management system, hereafter referred to as single environment, two different kernel regression models were used. The first model, which relies on a linear kernel, was the GBLUP as it is one of the most popular methods for genomic prediction (Van Raden 2008). For this model, the kernel matrix (*K*) was computed as *K = XX’, X* being the centered genotype matrix (-1, 0, 1) with N×P dimension, where N is the number of genotypes and P the number of markers. The second model, which is based on reproducing kernel Hilbert space (RKHS) approaches, used a Gaussian kernel *K(x_i_,x_j_) = exp(—h* || *x_i_ — x_j_||^2^*) to build the kernel matrix between the marker genotype vectors *x*_*i*_ and *x*_*j*_, where (*i,j) ϵ {1,…, N}*^*2*^. The bandwidth parameter *h* was estimated using the method described by Pérez-Elizalde et al. (2015) based on a Bayesian method that relies on the estimation of the mode of the joint posterior distribution of *h* and a form parameter φ. We used the R function *margh.fun* provided by Pérez-Elizalde et al. (2015) with a gamma prior distribution for h, with a shape parameter equal to 3, and a scale parameter equal to 1.5.

#### Multi-environment models

To predict the genomic estimated breeding values with data from the two water management systems, hereafter referred to as multi-environment prediction, we used the statistical models described above with extensions that integrate environmental effects. In the extended GBLUP model, the effects of *m* environments, and the effects of the P markers are separated into two components: the main effect of the markers for all the environments and the effect of the markers for each environment (Lopez-Cruz et al. 2015). For RKHS, we used two extended models incorporating G×E: RKHS-1 corresponding to the “Empirical Bayesian–Genotype × Environment Interaction Model” proposed by Cuevas et al. (2016), and RKHS-2 corresponding to the environmental model (3) proposed by Cuevas et al. (2017). Like the extended GBLUP, the first model (RKHS-1) considers the effects of *m* environments, and the effects of the markers are separated into a main effect for all the environments and an effect specific to each environment:

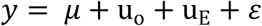

In this mixed model, *y* is the response vector, *μ* is the overall intercept, u_o_ captures the marker information among environments, and u_E_ accounts for the marker information in each environment. The random effects u_o_ follow a multivariate normal distribution with mean zero and a variance-covariance matrix 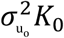, *K*_*0*_ constructed with the Gaussian kernel from the marker matrix *X*_*0*_.

The latter model (RKHS-2), considers that the performances of accessions in different environments are correlated such that there is a genetic correlation between environments that can be modeled with matrices of order m×m, *m* being the number of environments:

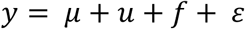

In this mixed model, y is the response vector, *μ* is the vector with the intercept of each environment, *u* the random vector of individual genetic values, *f* the genetic effects associated with individuals that were not accounted for in component *u*, and *£* the random vector of the error. *u, f* and *ε* are independent and normally distributed. For more methodological details concerning the extended GBLUP, RKHS-1 and RKHS-2 statistical models please refer to Lopez-Cruz et al. (2015), Cuevas et al. (2016) and to Cuevas et al. (2017), respectively.

#### Implementation of the models

Analyses were performed in the R 3.4.2 environment (R Core Team 2017) with the R packages *BGLR* 1.0.5 (Pérez and de los Campos 2014) and *MTM* 1.0.0 (De los Campos and Grüneberg 2018). For both packages, 35,000 iterations for the Gibbs sampler were used. For the inference, 3,000 samples were used after removing the first 5,000 samples (burn-in) and keeping one in ten samples to avoid auto-correlation (thinning). Convergence of Markov chain Monte Carlo algorithm was assessed for all parameters of the models with Gelman-Rubin tests (Gelman and Rubin 1992) using the R-package *coda* 0.19-1 (Plummer et al. 2006).

### Assessing genomic prediction accuracy

Prediction accuracy for the three traits and their related response to water management (index and slope) were assessed with two different validation schemes. The first scheme used only the RP with random partitions and is referred to hereafter as cross-validation. The second validation scheme used information from the RP to predict the performance of the PP (referred as progeny validation). The details of these two validation schemes are explained below.

#### Cross-validation within the reference population

Different types of random partitions were performed depending on the phenotypic and the genotypic information used in the statistical model. For traits in a single environment and for response variables, 80% of the 284 accessions (i.e. 227 accessions) of the RP were used as the training set and the remaining 20% (57 accessions) was used as the validation set. For multi-environment models, two different methods of cross-validation were applied. The first method (M1), which resembled what was done in the single environment, used 80% of the observations as a training set and the remaining 20% as the validation set and assumed that phenotypic observations for the two environments are available for the individuals composing the training set while no phenotypic data are available for the individuals in the validation set. M1 corresponds to the situation when the phenotypes of newly generated individuals have to be predicted based only on their genotypic information (Burgueño et al., 2012). The second method (M2) also used 80% of the observations as a training set and the remaining 20% as the validation set but assumed that at least one observation in one environment was available for the individuals in both the training set and the validation set. M2 corresponds to the situation when phenotypes in one environment have to be predicted with genotypic information and phenotypes from the other environment (Burgueño et al. 2012).

One hundred replicates were computed for all random partitioning in the training and validation sets. The prediction accuracy of each partition was calculated as the Pearson correlation coefficient between predictions and phenotypes in the validation set. For multi-environment models, the correlation was calculated within each environment. For each trait (FL, NI and PW) and each statistical model (GBLUP, RKHS-1 and RKHS-2), the same partitions were used to compute the prediction accuracy. The resulting estimates of prediction accuracy were averaged and the associated standard error was calculated.

To analyze the sources of variation of the prediction accuracy, the accuracy (*r*) of each prediction experiment was transformed into a Z-statistic using the equation: *Z =* 0.5 [ln(1 *+ r) —* ln(1 - *r)]* and used as a dependent variable in an analysis of variance. A separate analysis was performed for each trait. After estimating the confidence limits and means for Z, these were transformed back to *r* variables.

#### Progeny validation across populations

For progeny validation, the model was trained on the RP in order to predict the performance of the PP based on genotypic information. Three validation scenarios were evaluated. In the first scenario (S1) only the 31 parental lines were used as the training set. In the second scenario (S2), the CDmean method (Rincent et al. 2012) was used to select 100 accessions in the RP for the training set. In the third scenario (S3), all the RP accessions were included in the training set. In all three scenarios, the validation set was made up of all the PP lines. Like for cross-validation, prediction accuracy was calculated as the Pearson correlation coefficient between predictions and phenotypes in the validation set.

### Data Availability

The genotypic and phenotypic data are available in the TropGene database in the tab “Studies” as “GS-Ruse”: To access the TropGene database go to http://tropgenedb.cirad.fr/tropgene/JSP/interface.jsp?module=RICE).

## Results

### Analysis of the phenotypic variations and responses to water management

The partitioning of the observed phenotypic variation into different sources of variation via the mixed model analysis is shown in Table S1. Models were adjusted separately for each population (RP and PP) and each water management system (CF and AWD). Conditional R^2^ ranged from 0.33 to 0.96, indicating moderate to good fit of the model (Table 1). The lowest R^2^ values were obtained for NI trait in both populations and both conditions. The highest R2 values were obtained for FL. Whatever the trait or water management system considered, the genotype contributed significantly to the phenotypic variation in each population. A higher contribution of the genotype effect to the phenotypic variation was observed for FL compared to NI and to a lesser extent to PW. Broad-sense heritability (*H*^*2*^) tended to confirm this trend (Table 1). Indeed, depending on the population and the condition, *H*^*2*^ ranged from 0.85 to 0.94 for FL, from 0.75 to 0.90 for PW, and from 0.56 to 0.77 for NI. A slight increase in *H*^*2*^ was observed in CF water management compared to in AWD for FL and PW in RP. There was no significant difference in PP.

**Table 1:**
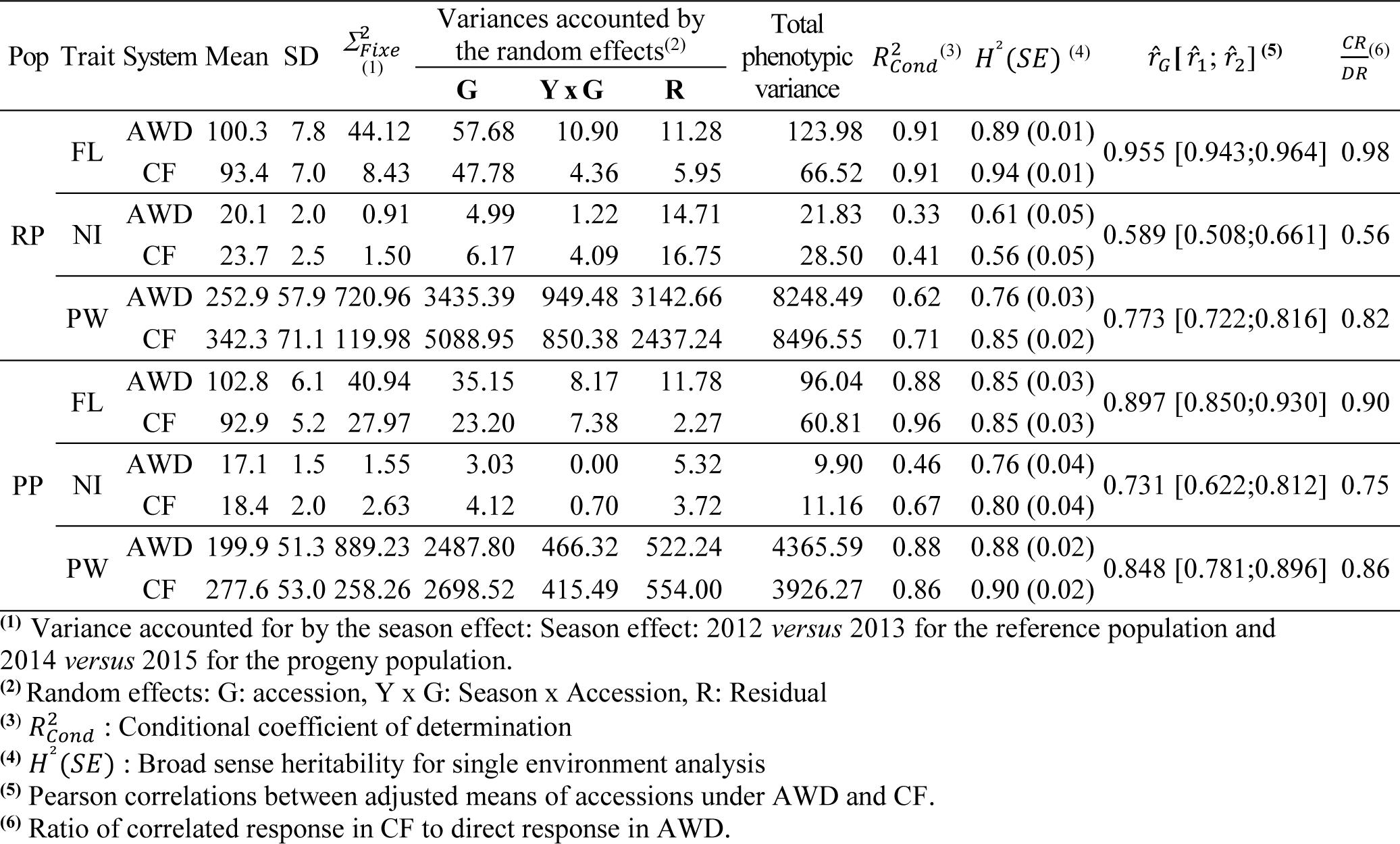
Sources of phenotypic variation and derived summary statistics of days to flowering (FL), nitrogen balance index (NI) and panicle weight (PW) in two populations of rice (reference RP and progeny PP) conducted in two consecutive seasons under two water management systems (AWD and CF).

| Pop | Trait | System | Mean | SD | $\Sigma_{Fixe}^2$<br>(1) | Variances accounted by<br>the random effects<sup>(2)</sup> | | | Total<br>phenotypic<br>variance | $R_{Cond}^2$<sup>(3)</sup> | $H^2(SE)$<sup>(4)</sup> | $\hat{r}_G[\hat{r}_1; \hat{r}_2]$<sup>(5)</sup> | $\frac{CR}{DR}$<sup>(6)</sup> |
| --- | --- | --- | --- | --- | --- | --- | --- | --- | --- | --- | --- | --- | --- |
|  |  |  |  |  |  | G | Y x G | R |  |  |  |  |  |
| RP | FL | AWD | 100.3 | 7.8 | 44.12 | 57.68 | 10.90 | 11.28 | 123.98 | 0.91 | 0.89 (0.01) | 0.955 [0.943;0.964] | 0.98 |
|  |  | CF | 93.4 | 7.0 | 8.43 | 47.78 | 4.36 | 5.95 | 66.52 | 0.91 | 0.94 (0.01) |  |  |
|  | NI | AWD | 20.1 | 2.0 | 0.91 | 4.99 | 1.22 | 14.71 | 21.83 | 0.33 | 0.61 (0.05) | 0.589 [0.508;0.661] | 0.56 |
|  |  | CF | 23.7 | 2.5 | 1.50 | 6.17 | 4.09 | 16.75 | 28.50 | 0.41 | 0.56 (0.05) |  |  |
|  | PW | AWD | 252.9 | 57.9 | 720.96 | 3435.39 | 949.48 | 3142.66 | 8248.49 | 0.62 | 0.76 (0.03) | 0.773 [0.722;0.816] | 0.82 |
|  |  | CF | 342.3 | 71.1 | 119.98 | 5088.95 | 850.38 | 2437.24 | 8496.55 | 0.71 | 0.85 (0.02) |  |  |
| PP | FL | AWD | 102.8 | 6.1 | 40.94 | 35.15 | 8.17 | 11.78 | 96.04 | 0.88 | 0.85 (0.03) | 0.897 [0.850;0.930] | 0.90 |
|  |  | CF | 92.9 | 5.2 | 27.97 | 23.20 | 7.38 | 2.27 | 60.81 | 0.96 | 0.85 (0.03) |  |  |
|  | NI | AWD | 17.1 | 1.5 | 1.55 | 3.03 | 0.00 | 5.32 | 9.90 | 0.46 | 0.76 (0.04) | 0.731 [0.622;0.812] | 0.75 |
|  |  | CF | 18.4 | 2.0 | 2.63 | 4.12 | 0.70 | 3.72 | 11.16 | 0.67 | 0.80 (0.04) |  |  |
|  | PW | AWD | 199.9 | 51.3 | 889.23 | 2487.80 | 466.32 | 522.24 | 4365.59 | 0.88 | 0.88 (0.02) | 0.848 [0.781;0.896] | 0.86 |
|  |  | CF | 277.6 | 53.0 | 258.26 | 2698.52 | 415.49 | 554.00 | 3926.27 | 0.86 | 0.90 (0.02) |  |  |

<sup>(1)</sup> Variance accounted for by the season effect: Season effect: 2012 *versus* 2013 for the reference population and 2014 *versus* 2015 for the progeny population.
<sup>(2)</sup> Random effects: G: accession, Y x G: Season x Accession, R: Residual
<sup>(3)</sup> $R_{Cond}^2$ : Conditional coefficient of determination
<sup>(4)</sup> $H^2(SE)$ : Broad sense heritability for single environment analysis
<sup>(5)</sup> Pearson correlations between adjusted means of accessions under AWD and CF.
<sup>(6)</sup> Ratio of correlated response in CF to direct response in AWD.

The three traits investigated exhibited normal distribution in the RP and PP under both AWD and FC (Figure 1). The AWD water management resulted in medium intensity stress for FL (7.4% and 10.8% for RP and PP, respectively) and NI (−15.6% and −7.6%), and in rather severe stress intensity for PW (-26.6% and −27.9%). On average, both populations flowered significantly later under AWD than CF. The average FL values were 100.3 (102.8) in AWD and 93.4 (92.9) in CF, for RP and (PP). Conversely, significantly lower NI and PW values were observed in AWD compared to CF in both populations. For PW, the average differences between the two water management systems were 89.4 g for RP and 77.7 g for PP. For NI, in addition to differences in the average performance of the two water management systems, significant differences in distribution were also observed between RP and PP, for the extent of diversity, much larger for the RP, and for the frequency of individuals with low NI, much higher in the PP (Figure S1).

**Figure 1:**
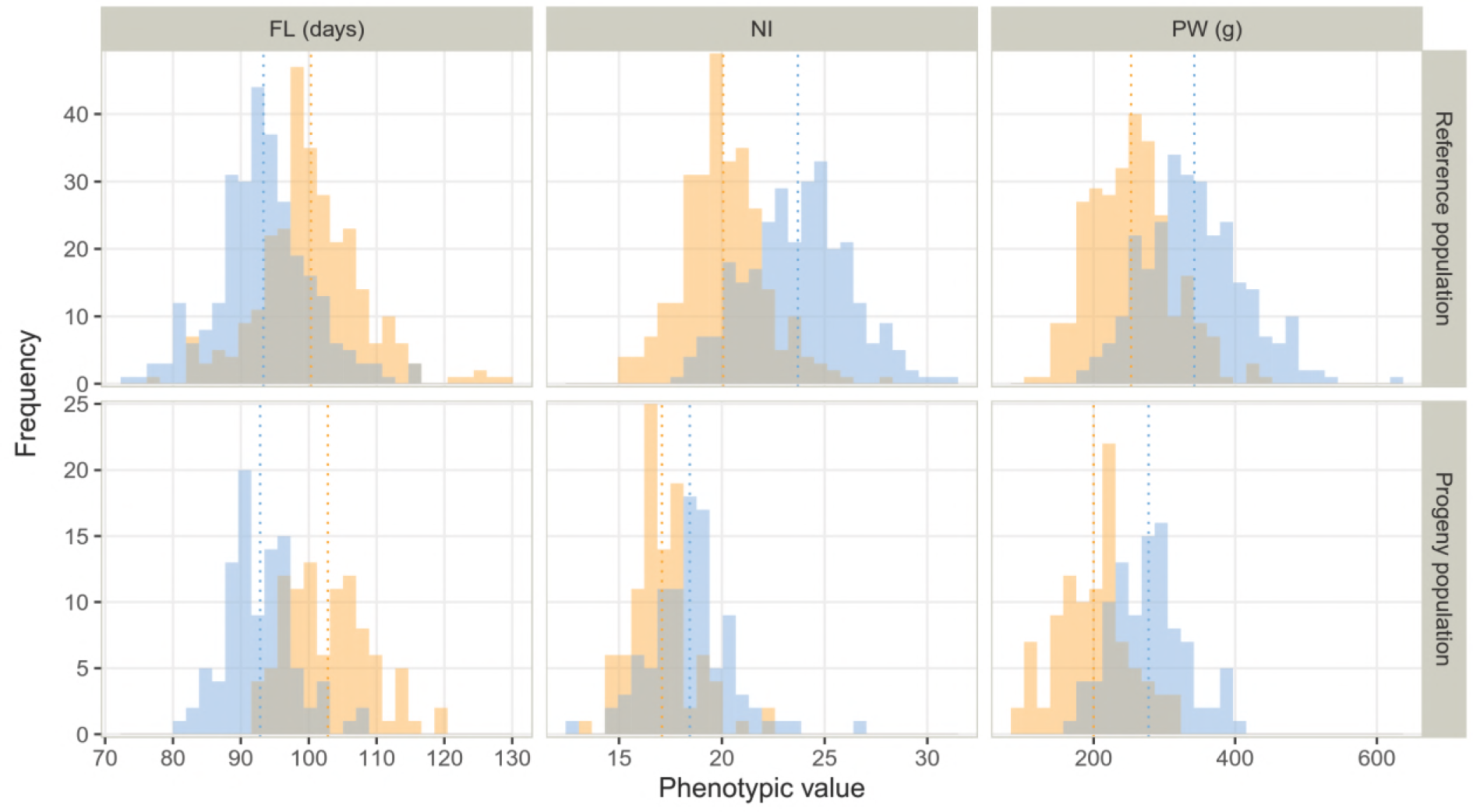
Distribution of adjusted phenotypic values of days to flowering (FL), nitrogen balance index (NI) and panicle weight (PW) within the reference and progeny populations in continuous flooding (blue) and alternate wetting and drying (orange) conditions.

Partitioning of the phenotypic variation from the two water management systems into different sources of variation revealed the existence of significant interactions between genotypes and water management systems in both RP and PP, for all traits except FL in RP (Table S2). For all traits and populations, the ranking of the individuals was affected by water management and the Spearman’s rank correlation coefficients between traits values under the two water management systems were medium to high (Figure 2). As a result, for each trait the ratio of correlated response to selection under FC, relative to direct response to selection under AWD, ranged from medium (0.56 and 0.75 for NI) to very high (0.98 and 0.90 for FL), suggesting indirect selection for adaptation to AWD is feasible (Table 1).

**Figure 2:**
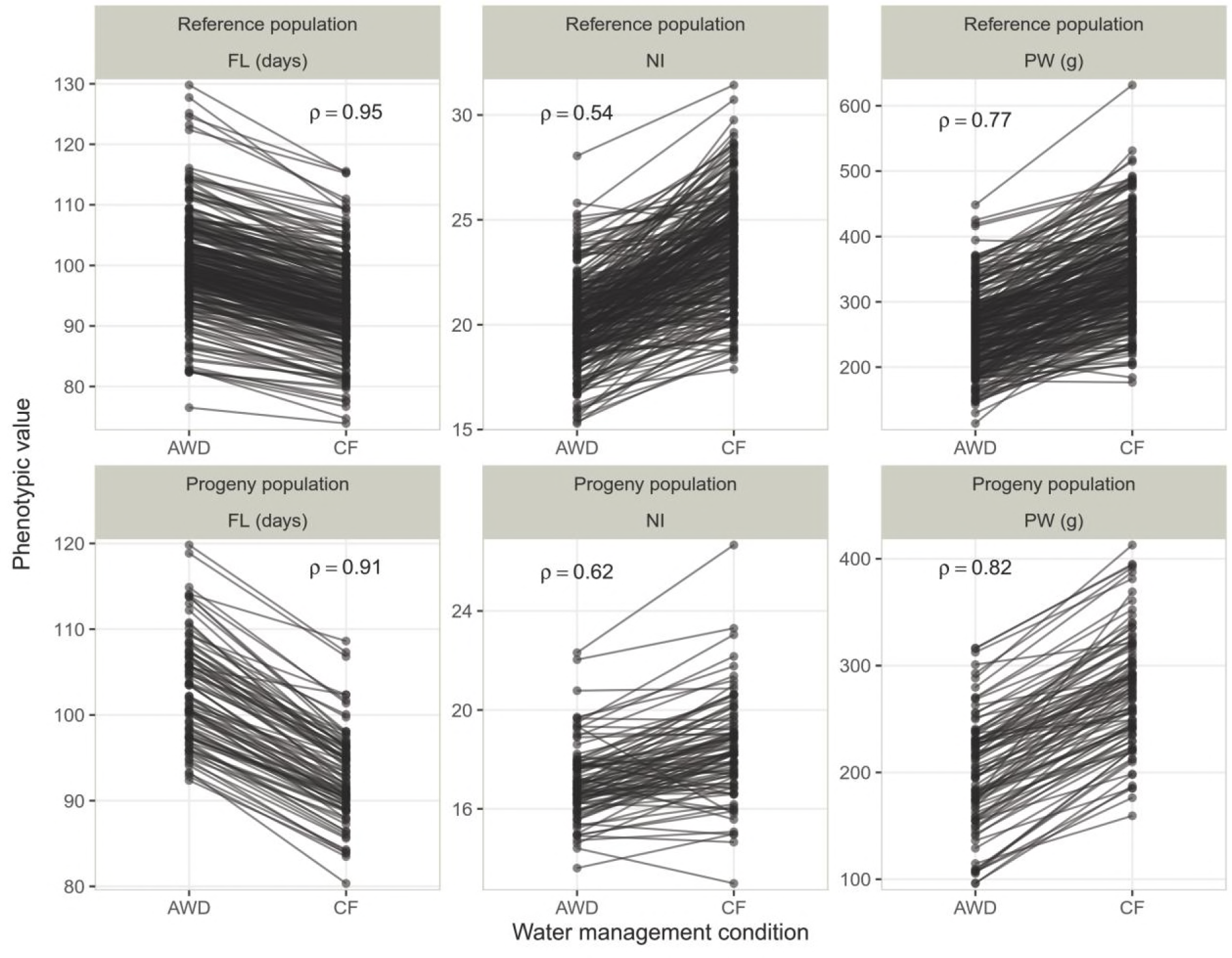
Reaction norm between the two conditions (AWD: alternate wetting and drying and CF: continuous flooding) for all the genotypes of the two populations (the reference population and the progeny population). The three traits are represented: days to flowering (FL), nitrogen balance index (NI) and panicle weight (PW). Spearman’s rank correlation coefficient (ρ) is indicated in each panel.

The two computed variables (index and slope) characterizing the accessions’ response to AWD, revealed a Gaussian distribution for the three phenotypic traits considered (Figure S1). An ANOVA of these computed variables revealed significant genotype effects on the three traits in both RP and PP populations (Table S3). By construction, the correlations between phenotypic values per condition and the slope were higher than those with the index whatever the trait and the population considered. Interestingly, the index behaved differently in each trait (Figure S1). For FL, low correlations were found either with AWD or CF. For NI, higher correlations were found with CF (- 0.51 for RP and - 0.58 for PP) than with AWD (0.39 for RP and 0.13 for PP). For PW, correlations were higher with AWD (0.42 for RP and 0.71 for PP) than with CF (- 0.23 for RP and 0.24 for PP). For the three traits considered, there was almost no correlation between the index and the slope variables (Figure S1): FL (0.12 for RP and 0.17 for PP), NI (0.-0.16 for RP and −0.31 for PP) and PW (-0.03 for RP and 0.04 for PP).

### Accuracy of genomic prediction for the response variables

#### Prediction accuracy in the reference population

The average prediction accuracies obtained for the two response variables were compared with those obtained for the observed variables in each water management system considered as references (Table 2). The overall mean accuracy for the observed variables (the three traits under the two water management systems), and for the response variables, was 0.54 but the range extended from −0.12 to 0.88, depending on the prediction model, the trait and the type of variable (**Erreur ! Source du renvoi introuvable.**, Table S4). The most significant factor influencing accuracy was the type of variable (Table 2). Indeed, regardless of the trait or the statistical model, accuracy for the index was lower than for the slope: 0.31 against versus 0.64 on average (**Erreur ! Source du renvoi introuvable.**). Interestingly, NI, which presented the highest G×E, was the trait with the lowest accuracy for the index (0.17 and 0.21). However, index predictions were less accurate for FL, the trait with the lowest G×E, (0.29 and 0.30) than for PW (0.43 and 0.48) with intermediate G×E. In agreement with the medium to high correlations at phenotypic level, the prediction accuracies for the slope and the variables under each condition were comparable. However, different trends were observed depending on the trait. For FL and PW, accuracies for the slope were closer to accuracies under AWD than under CF. For NI, the opposite was observed. In all cases, slope prediction was as accurate as the best single-environment prediction. The level of accuracy depended secondly on the trait (Table 2). On average, accuracy was higher for FL (0.6) than for PW (0.58) and NI (0.45). The statistical models differed significantly from each other although the effect was small. RKHS performed better than GBLUP in almost all cases with differences in accuracy of up to 0.05. The interactions between factors influencing prediction accuracy were not important, except for the one between the response variable and the trait (Table 2).

**Table 2:** Analysis of factors that influence the prediction accuracy of response variables in the reference population. The effects of the type of response (index, slope, AWD, CF), the trait (FL, NI and PW), the statistical model (GBLUP and RKHS) and their interactions were evaluated.

| R<sup>2</sup> | CV | RMSE | Mean | Source | DF | SS | MS | FValue | ProbF |
| --- | --- | --- | --- | --- | --- | --- | --- | --- | --- |
| Model 1: Only main effects |  |  |  |  |  |  |  |  |  |
| 0.648 | 23.617 | 0.152 | 0.642 | Model | 6 | 101.489 | 16.915 | 734.86 | <0.0001 |
|  |  |  |  | Error | 2393 | 55.082 | 0.023 |  |  |
|  |  |  |  | Corrected Total | 2399 | 156.570 |  |  |  |
|  |  |  |  | Response | 3 | 77.532 | 25.844 | 1122.78 | <0.0001 |
|  |  |  |  | Trait | 2 | 23.571 | 11.785 | 512.01 | <0.0001 |
|  |  |  |  | S model | 1 | 0.386 | 0.386 | 16.76 | <0.0001 |
| Model 2: Main effects and interactions |  |  |  |  |  |  |  |  |  |
| 0.732 | 20.681 | 0.133 | 0.642 | Model | 23 | 114.633 | 4.984 | 282.38 | <0.0001 |
|  |  |  |  | Error | 2376 | 41.937 | 0.018 |  |  |
|  |  |  |  | Corrected Total | 2399 | 156.570 |  |  |  |
|  |  |  |  | Response | 3 | 77.532 | 25.844 | 1464.21 | <0.0001 |
|  |  |  |  | Trait | 2 | 23.571 | 11.785 | 667.71 | <0.0001 |
|  |  |  |  | S model | 1 | 0.386 | 0.386 | 21.86 | <0.0001 |
|  |  |  |  | Response*Trait | 6 | 12.456 | 2.076 | 117.61 | <0.0001 |
|  |  |  |  | Trait*S model | 2 | 0.433 | 0.217 | 12.27 | <0.0001 |
|  |  |  |  | Response*S model | 3 | 0.073 | 0.024 | 1.38 | 0.2459 |
|  |  |  |  | Response*Trait*S model | 6 | 0.182 | 0.030 | 1.72 | 0.1126 |

R<sup>2</sup>: Coefficient of determination; CV: Coefficient of variation; RMSE: Root mean square error; Mean: Intercept value of the transformed accuracy (Z); DF: Degree of freedom; SS: Sum of squares; MS: Mean square.

#### Prediction accuracy across populations

On average, across generation prediction for both observed and computed response variables was less accurate (0.28) than prediction within the reference population (Table S5). Accuracies ranged from −0.01 to 0.38, with an average of 0.25 for FL, from −0.1 to 0.45, with an average of 0.22, for NI, and from 0.14 to 0.56, with an average of 0.38 for PW, depending on the type of variables (observed variables, index and slope), the scenario and the model (Figure 4). Among these factors, the most influential was again the type of response (Table S5), with the lowest average accuracy of 0.12 for index and the highest average accuracy of 0.35 for slope. The prediction accuracy under the single environment AWD and CF averaged 0.34 and 0.32, respectively. The effect of the scenario came in second, with an average accuracy of 0.27 for S1, 0.22 for S2, and 0.35 for S3. The statistical models GBLUP and RKHS performed similarly on average (accuracy of 0.28) but the range of variation was slightly wider in RKHS (-0.1 to 0.56) than in GBLUP (-0.01 to 0.51).

**Figure 3:**
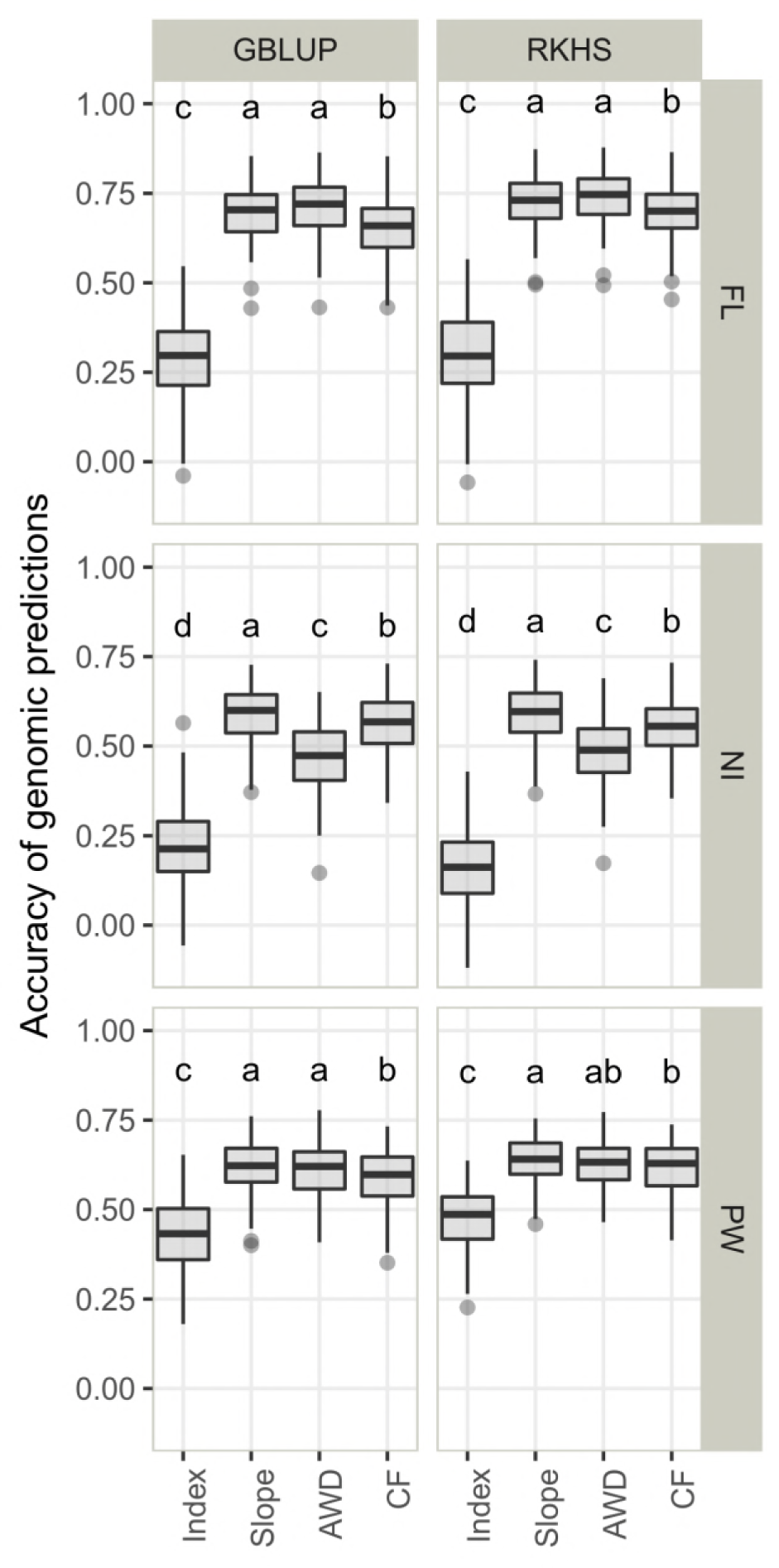
Accuracy of genomic prediction in cross validation experiments within the reference population obtained with two statistical models (GBLUP, RKHS) for the response variables (index and slope) and the performance within each condition (AWD and CF). The three traits are presented: days to flowering (FL), nitrogen balance index (NI) and panicle weight (PW). The letters in each panel represent the results of Tukey’s HSD comparison of means and apply to each panel independently. The means differ significantly (p-value < 0 05) if two boxplots have no letter in common.

**Figure 4:**
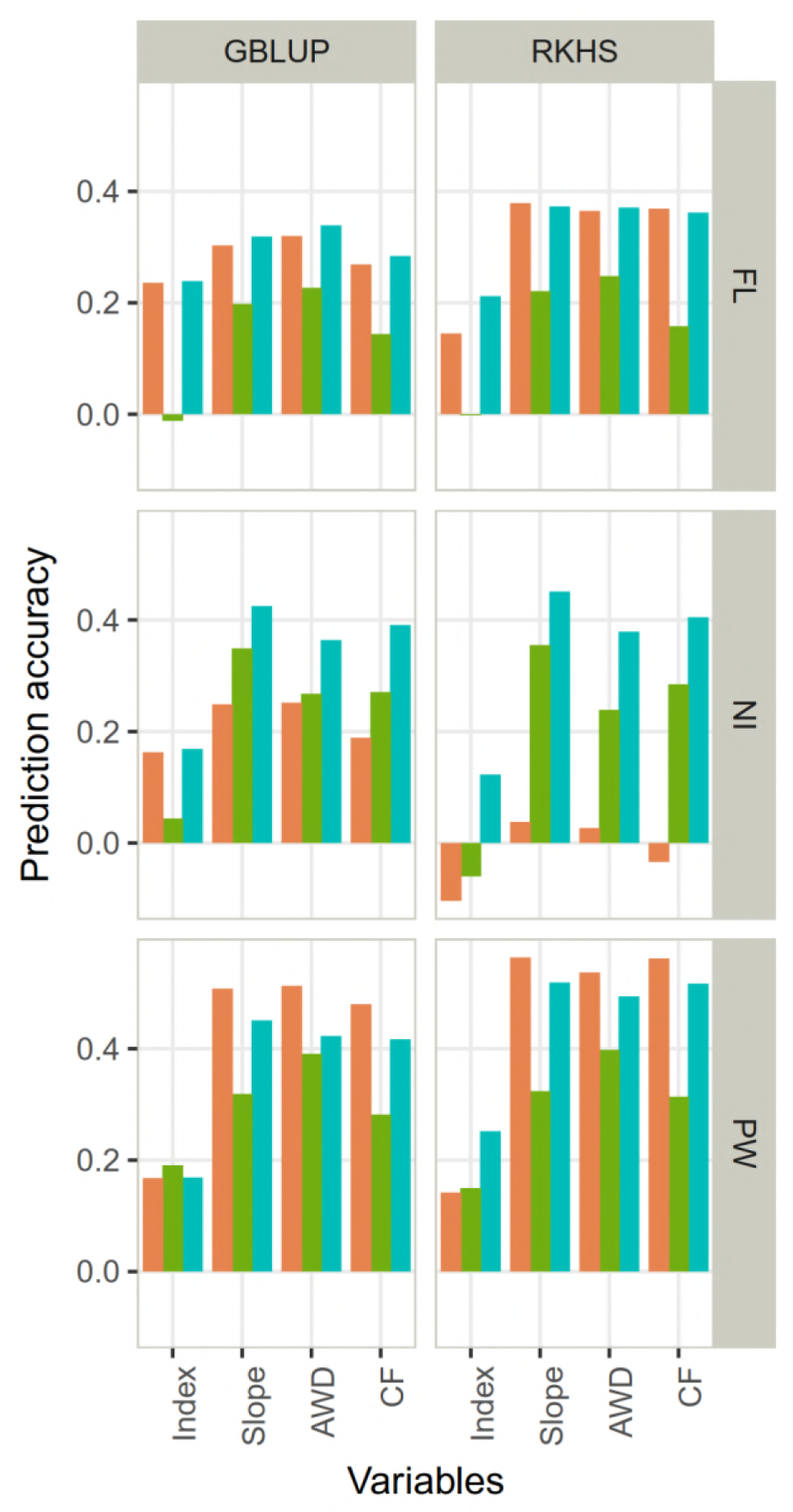
Accuracy of genomic prediction in across population validation for the response variables (index and slope) and the performance within each condition (AWD and CF) obtained. Two statistical models (GBLUP, RKHS) and three traits (days to flowering (FL), nitrogen balance index (NI) and 100 panicle weight (PW)) were studied. The scenarios used to define the training set are in color: orange (S1: only the parents), green (S2: 100 individuals of the RP selected with CDmean) and blue (S3: the whole RP).

### Accuracy of genomic prediction using multi-environment models

#### Prediction accuracy in the reference population

The focus here was on multi-environment models and the two different cross-validation methods (M1 and M2), using single environment models as the baseline. Average accuracies ranged from 0.47 to 0.96, depending on (in decreasing importance): the trait, the type of model (i.e. single versus multi-environment), the cross-validation strategy, the statistical model and the water management system (Figure 5, Table S6). The average accuracy was of 0.79, 0.56 and 0.69 for FL, NI and PW respectively. Whatever the trait or the water management system, multi-environment models with the M1 strategy performed similarly to the single environment model with a decrease of up to 0.02 for GBLUP and up to 0.03 for RKHS-1 and RKHS-2. As expected, the multi-environment models with the M2 strategy outperformed single environment models with an average gain of 0.23 and 0.27 for FL, 0.14 and 0.10 for NI and 0.20 and 0.20 for PW in AWD and CF, respectively. These gains in accuracy were in agreement with the level of G×E found for each trait. Among the significant interactions between factors, the trait × cross validation strategy interaction was the most important and corresponded to a scale interaction (Table 3). Among the multi-environment prediction models, RKHS-1 and RKHS-2 performed similarly, with average accuracy of 0.72 and 0.71, respectively, and performed systematically slightly better than GBLUP, with a gain in accuracy of up to 0.04.

**Figure 5:**
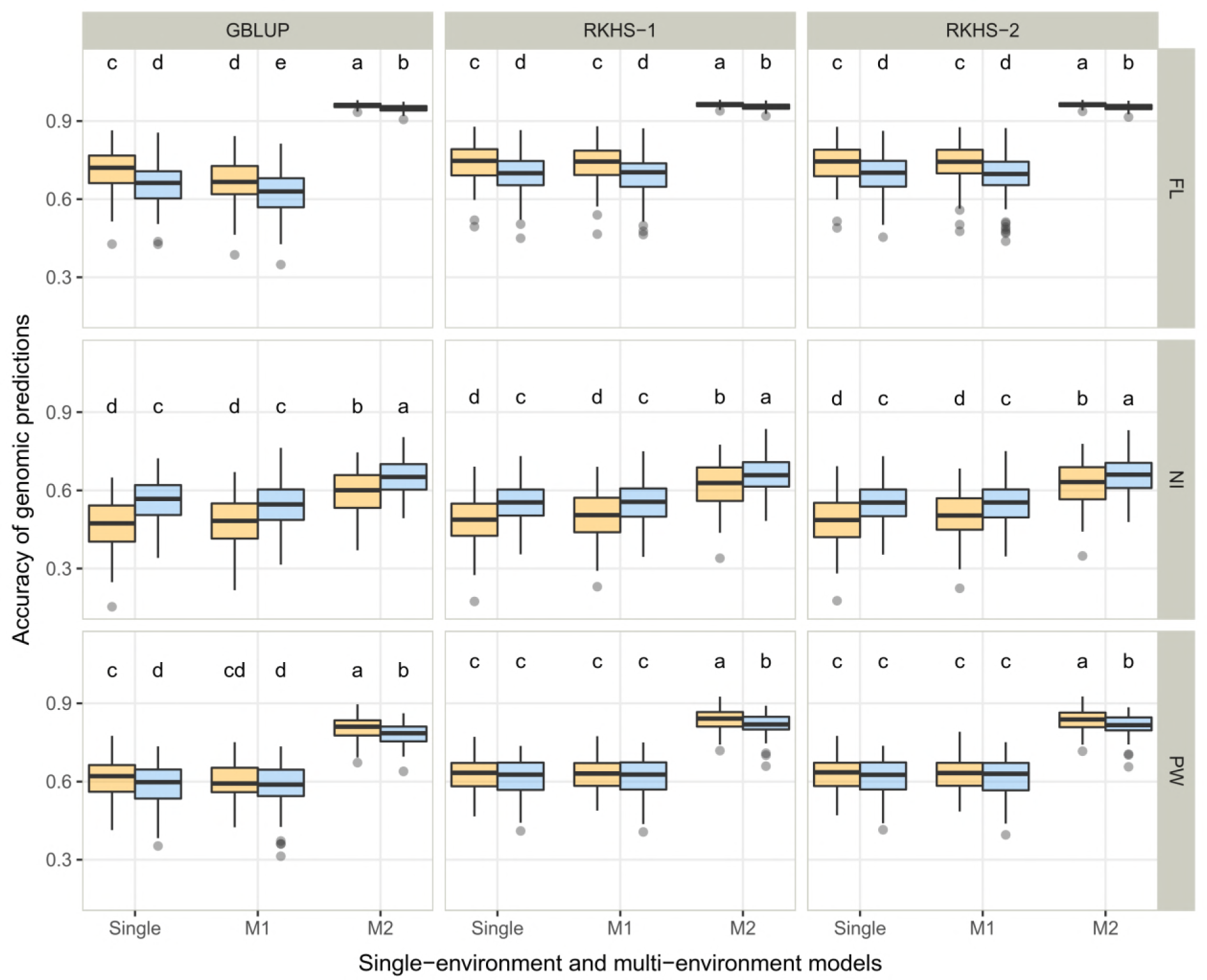
Single environment and multi-environment (M1 and M2) prediction accuracy in cross validation experiments in the reference population obtained with three statistical models (GBLUP, RKHS-1, RKHS-2). Continuous flooding and alternate wetting and drying water management conditions are in blue and orange, respectively. The three studied traits are presented: days to flowering (FL), nitrogen balance index (NI) and panicle weight (PW). The letters in each panel represent the results of Tukey’s HSD comparison of means and apply to each panel independently. The means differ significantly (p-value < 0 05) if two boxplots have no letter in common.

**Table 3:** Analysis of factors that influence the variation in prediction accuracy in the reference population using multi-environment models. The effects of the statistical model (GBLUP, RKHS-1 and RKHS-2), the trait (FL, NI and PW), the cross-validation strategy (M1 and M2) and the target condition (AWD and CF) and their interactions were evaluated.

| R<sup>2</sup> | CV | RMSE | Mean | Source | DF | SS | MS | FValue | ProbF |
| --- | --- | --- | --- | --- | --- | --- | --- | --- | --- |
| Analysis with only main effects |  |  |  |  |  |  |  |  |  |
| 0.723 | 24.163 | 0.221 | 0.914 | Model | 7 | 687.496 | 98.214 | 2014.66 | <0.0001 |
|  |  |  |  | Error | 5392 | 262.858 | 0.049 |  |  |
|  |  |  |  | Corrected Total | 5399 | 950.354 |  |  |  |
|  |  |  |  | CV strategy | 2 | 362.879 | 181.439 | 3721.86 | <.0001 |
|  |  |  |  | Trait | 2 | 320.946 | 160.473 | 3291.78 | <.0001 |
|  |  |  |  | S model | 2 | 3.352 | 1.676 | 34.38 | <.0001 |
|  |  |  |  | Target condition | 1 | 0.319 | 0.319 | 6.55 | 0.0105 |
| Analysis with main effects and all first-order interactions |  |  |  |  |  |  |  |  |  |
| 0.899 | 14.640 | 0.134 | 0.914 | Model | 25 | 854.176 | 34.167 | 1909.11 | <.0001 |
|  |  |  |  | Error | 5374 | 96.178 | 0.018 |  |  |
|  |  |  |  | Corrected Total | 5399 | 950.354 |  |  |  |
|  |  |  |  | CV strategy | 2 | 362.879 | 181.440 | 10138.0 | <.0001 |
|  |  |  |  | Trait | 2 | 320.946 | 160.473 | 8966.54 | <.0001 |
|  |  |  |  | S model | 2 | 3.352 | 1.676 | 93.65 | <.0001 |
|  |  |  |  | Target condition | 1 | 0.319 | 0.319 | 17.83 | <.0001 |
|  |  |  |  | CV strategy*Trait | 4 | 157.483 | 39.371 | 2199.87 | <.0001 |
|  |  |  |  | Target condition*Trait | 2 | 7.811 | 3.906 | 218.23 | <.0001 |
|  |  |  |  | Trait*S model | 4 | 0.783 | 0.196 | 10.94 | <.0001 |
|  |  |  |  | Target condition*CV strategy | 2 | 0.300 | 0.150 | 8.37 | 0.0002 |
|  |  |  |  | CV strategy*S model | 4 | 0.300 | 0.075 | 4.20 | 0.0022 |
|  |  |  |  | Target condition*S model | 2 | 0.003 | 0.002 | 0.09 | 0.9169 |

R<sup>2</sup>: Coefficient of determination; CV: Coefficient of variation; RMSE: Root mean square error; Mean: Intercept value of the transformed accuracy (Z); DF: Degree of freedom; SS: Sum of squares; MS: Mean square.

#### Prediction accuracy across populations

The overall mean accuracy was 0.33, with values ranging from −0.03 up to 0.58 (Figure 6, Table S7), mainly depending on traits and scenarios for the composition of the training set. The average accuracy was of 0.30, 0.27, and 0.44 for FL, NI and PW, respectively. The average accuracy of the three scenarios was 0.32, 0.28 and 0.40 for S1, S2 and S3, respectively. The range of variation in accuracy for the remaining factors (single versus multiple environment, target environment and prediction model) did not exceed 0.03. These latter factors influenced the accuracy mainly in interactive mode.

**Figure 6:**
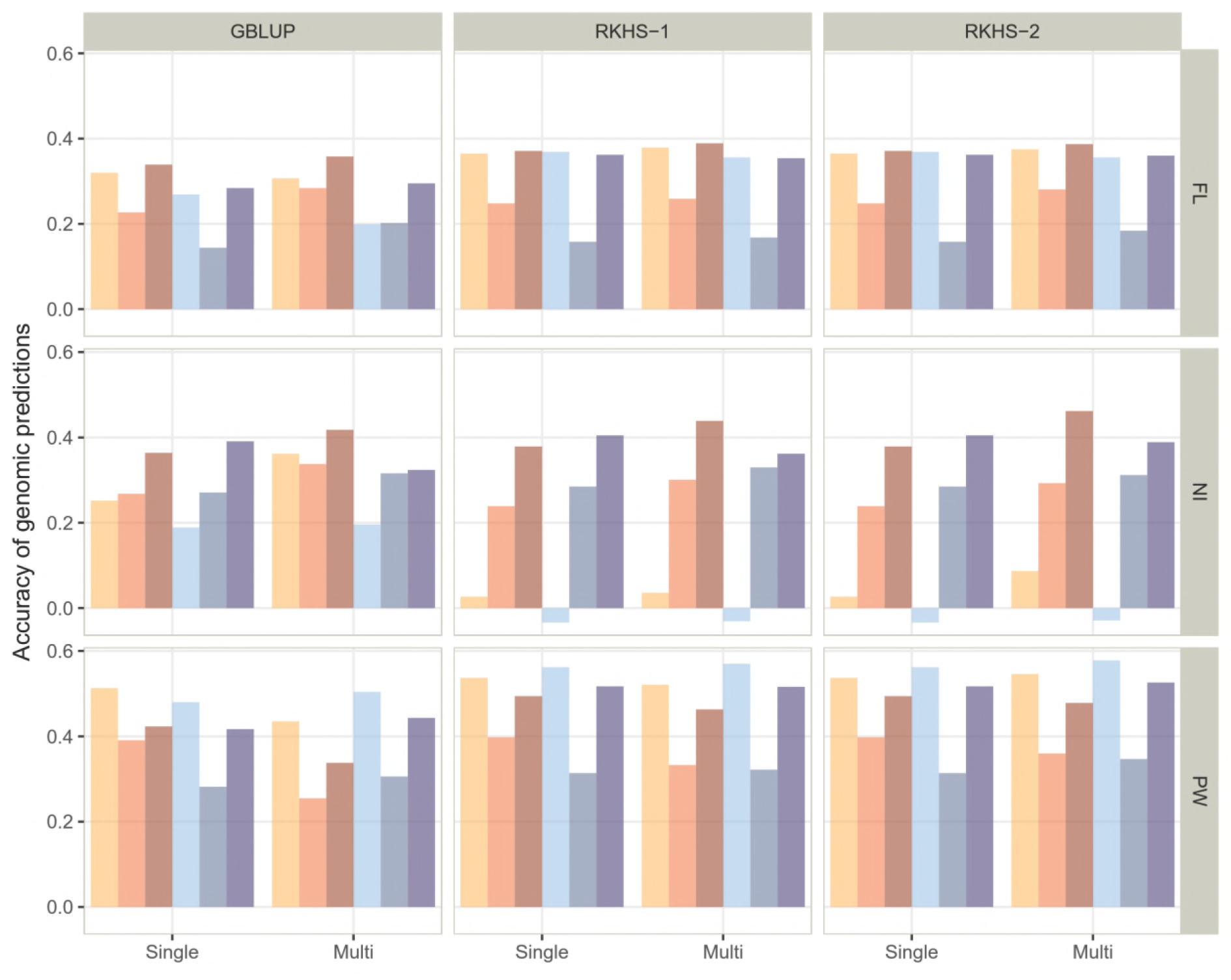
Single environment and multi-environment prediction accuracy in across population validation experiments obtained with three statistical models (GBLUP, RKHS-1, RKHS-2). Continuous flooding and alternate wetting and drying water management conditions are in blue and orange, respectively. The scenarios used to define the training set are represented by the different shades of orange or blue: light (S1: only the parents), intermediate (S2: 100 individuals of the RP selected with CDmean) and dark (S3: the whole RP). The three studied traits are presented: days to flowering (FL), nitrogen balance index (NI) and panicle weight (PW).

## Discussion

### Impact of AWD water management system on rice performance

The AWD water management implemented in this study (a new cycle of irrigation was triggered when soil water potential reached −30 kPa) resulted in medium intensity stress for FL and NI traits, rather severe stress intensity for PW when evaluated in terms of relative performance. The effects of AWD we observed on PW (-27% on average), are similar to those reported by Carrijo et al. (2017) on yield, in their review of 56 studies with 528 side-by-side comparisons of yield under AWD and CF. These authors reported an average decrease in yields of 5.4%, almost no yield losses under mild AWD (i.e. when soil water potential was kept ≥ −20 kPa), and yield losses of 22.6% relative to CF under severe AWD, when the soil water potential went beyond −20 kPa. However, in contrast with our experiment, which pioneered the analysis of genotypic responses to AWD within a diversity panel representing a large share of diversity of one of the sub-species of rice (*O. sativa, japonica*), the majority of the studies included in Carrijo et al.’s (2017) meta-analysis used only a small number of rice varieties and the crop was established by transplanting. Among the few studies reporting on traits other than grain yield, Sudhir et al. (2011) reported crop maturity delay of 5-10% under severe AWD, similar to our results (9% on average).

### Genomic prediction of response to AWD

The two computed variables (response index and slope of the joint regression) were intended to provide a measurement of G×E for each accession of RP and PP, which could be used as the entry phenotype for genomic prediction. The index, which evaluates tolerance to AWD water management, was very closely correlated with the stress sensitivity and tolerance index proposed by Fischer and Maurer (1978) and (Rosielle and Hamblin 1981), respectively (data not shown). The slope provides a measurement of stability of breeding material along environmental gradients in multi-environment trials (Eberhart and Russell 1966; Lin et al. 1986). However, the fact that the environmental index is not independent of the performances of the studied genotypes can introduce a bias in the estimate of the regression parameters (Crossa 1990). Moreover, the percentage of G×E variance explained is often very low, below 25% (for a review, see Brancourt-Hulmel et al. 1997). In our case, the number of environments considered, two, was probably too few for a precise estimate of the regression slope for each genotype. On the other hand, the large number of genotypes involved in the estimate of the environmental index (284 for RP and 97 for PP) limited the above-mentioned risk of bias. Given the very high correlations between the computed slopes and the measured phenotypes for the three traits under AWD and CF in both RP and PP populations (r > 0.9, except for PW under AWD in PP (r = 0.73), it represents a reasonably good single entry phenotype to consider for breeding both for adaptation to AWD and performance under CF.

The accuracy of genomic prediction for the response index was significantly lower than for the slope and for the corresponding measured traits under AWD and CF, suggesting limited genetic control of variation for the response index. Similar results were reported by (Huang et al. 2016) for trait stability in wheat. Nevertheless, given the loose correlations between the response index and the measured traits, genomic prediction for the index and the measured trait in CF could be used to select for good performance in both systems.

### Genomic prediction using multi-environment data

The potential of GS to accelerate the pace of genetic gains in major field crops has been documented by a large number of studies using a simulation approach or experimental data (Crossa et al. 2017; Hickey et al. 2017). In the case of rice, several empirical studies, summarized in Ben Hassen et al. (2017), confirmed this potential. However, the focus of most previous crop genomic prediction studies was on within-environment prediction, based on single environment models. It was recently demonstrated that the accuracy of genomic prediction models that account for G×E is significantly greater than that attained by single environment models (Cuevas et al. 2016; Cuevas et al. 2017; Burgueño et al. 2012; Jarquín et al. 2014; Lopez-Cruz et al. 2015; Heslot et al. 2014). The empirical component of almost all of these studies was based on data from unmanaged multi-environment trials of genotypes across several locations (and often several years), mainly conducted to study G×E and the general stability of the genotype across environments. The multi-environment genomic prediction results we present here stand out among the aforementioned ones because we used data from managed bi-environment trials undertaken to study G×E and genotype adaptation to a specific abiotic constraint, i.e. AWD water management.

The level of prediction accuracy obtained in our cross validation experiments in the reference population under the M1 prediction strategy with the multi-environment GBLUP, RKHS-1 and RKHS-2 models, calibrated with data from both AWD and CF water management, was similar to that obtained with their single environment counterparts, calibrated with data from either AWD or CF. These results confirm the power of multi-environment genomic models to predict the performances of untested genotypes using data from multiple trials. Under the M2 prediction strategy, the three multi-environment models provided significantly higher prediction accuracy for genotypes that had not been tested in one of the two water management systems than their single-environment counterparts, further confirming the advantages of multi-environment prediction models. In order to challenge the performance of the multi-environment models further, we ran the M1 and M2 strategies with a larger number of untested entrees (40% instead of 20%) in both AWD and CF for M1, in AWD or CF for M2. The results in Figure S2 show a very small reduction in prediction accuracy. The average prediction accuracy for the three traits, the two water managements and the three prediction models was 0.59 instead of 0.61 for M1, and 0.79 instead of 0.81 for M2. These results suggest the possibility of optimizing the method of evaluation of the lines by targeting a specific set of lines for each condition (Rincent et al. 2017).

Lopez-Cruz et al. (2015) reported gains in prediction accuracy of up to 30% with the GBLUP-type multi-environment model compared to an across-environment analysis that ignores G×E, when applied to the wheat grain yield of three sets of advanced lines recorded in three different years under three irrigation regimes. In our case, significant gains in accuracy were observed only with the M2 strategy, and ranged from 17% for NI to 29% for FL. Using wheat and maize data, Cuevas et al. (2016) reported up to 68% higher accuracy for RKHS-1 models compared to single environment models and up to 17% compared to GBLUP-G×E. These authors hypothesized that the superiority of the Gaussian kernel models over the linear kernel was due to more flexible kernels that account for small, more complex marker main effects and marker specific interaction effects. In our experiments, RKHS-1 was up to 35% more accurate than single environment GBLUP and up to 10% more accurate than GBLUP-G×E model. On the other hand, we did not observe any notable differences in the prediction accuracy of the RKHS-2 model compared to GBLUP-G×E and RKHS-1, as already reported by Cuevas et al. (2017). This is probably due to the positive correlation between performances under AWD and CF water management systems in our experiments, while the most favorable context for the application the approach developed by Cuevas et al. (2017) is said to be when different types of correlation (positive, zero, or negative) between the environments considered, coexist.

The results of our progeny validation experiments did not question the higher prediction accuracy of multi-environment models compared to single environment ones observed in our cross validation experiments in the reference population. However, in progeny validation experiments, the multi-environment models affected prediction accuracy mainly in interaction with other factors, such as the composition of the training set and the trait considered. These results also confirmed the important role of relatedness between the training and the validation set in prediction accuracy. It also confirmed the fact that relatively high accuracy could be achieved using only a rather small share of the RP, the most closely related to the PP as the training set, as reported by Ben Hassen et al. (2017).

Finally yet importantly, in both cross validation and progeny validation experiments, the multi-environment approach achieved higher prediction accuracy than the genomic prediction for the response index and the slope of the joint regression. For instance, compared to prediction for slope, the mean advantage of multi-environment prediction was 8% and 10% with GBLUP-G×E and RKHS-1 models, respectively. The advantage reached 25% under the M2 strategy of predicting unobserved phenotypes. In the progeny-validation experiments, the mean advantage was 20% and reached 30% under the S2 scenario of composition of the training set. To our knowledge, this finding has not yet been reported in the literature. It opens new perspectives in breeding for adaptation to AWD and to other abiotic stresses.

### Practical implications for breeding rice for adaptation to AWD

“More rice with less water” is vital for food security and for the sustainability of irrigated rice cropping systems (Tuong et al. 2005). AWD water management is one of the most widely used water-saving techniques practiced today (Carrijo et al. 2017). The development of rice varieties adapted to AWD, i.e. with as high yields as the best high yielding variety under CF, would greatly contribute to wider adoption of AWD water management by farmers (Price et al. 2013; Volante et al. 2017). Given the genetic diversity we observed for response to AWD within the working collection of the CREA, which represents only a share of the genetic diversity of the rice *japonica* sub-species, one can expect large genetic diversity at the whole species level.

The almost identical and high level of broad-sense heritability observed under AWD and CF water management systems demonstrates the feasibility of direct selection for AWD. Such high heritability under managed abiotic stress has already been reported in rice for grain yield under drought stress (Venuprasad et al. 2007; Kumar et al. 2008). However, the adoption of the direct selection option may not be practicable for breeding programs with limited resources, if they also need to continue to breed for CF water management. Moreover, this option would not take full advantage of historical data produced by the breeding program for CF. The high accuracy of multi-environment genomic prediction we observed in the present study, especially in across-environment prediction, paves the way for a new breeding option: conducting simultaneously direct and indirect selection for both AWD and CF. Indeed, as we saw in our M2 strategy, the multi-environment genomic models can boost the predictive power of across-environment predictions, i.e. from CF to AWD and *vice versa.* In this context, the practical question would be the number of selection candidates that need to be phenotyped under the two water management systems relative to the number of candidates that need to be phenotyped under one water management system only. Our results suggest that, for the germplasm and environmental conditions we used and the traits we considered, the percentage of untested candidates under AWD can go up to 40% with no significant negative effect on prediction accuracy as long as they are tested under CF, or *vice versa*. Considering the additional cost reductions that could be obtained by optimizing the size of the training set, as shown by the S1 scenario in our across-generations prediction experiments, it seems possible to add the objective of adaptation to AWD to an existing GS based rice breeding program for CF, with rather limited additional costs. Ben Hassen et al. (2017) showed that rice breeding programs based on pedigree schemes can use a genomic model trained with data from their working collection to predict performances of progenies produced by the conventional pedigree breeding program. Breeding for adaptation to AWD can be integrated in this general scheme. The feasibility of application of this breeding approach to other abiotic stresses deserves further exploration.

## Authors’ contributions

NA and GV conceived the study. JB, MBH and TVC analyzed the data. MBH, TVC, JB and NA wrote the manuscript.

## Acknowledgments

This work was funded by Agropolis Foundation (http://www.agropolis-fondation.fr/) and Cariplo Foundation (http://www.fondazionecariplo.it/), Grant no 1201-006.

## Conflict of Interest

The authors declare that they have no conflict of interest.

## supplementary figures and tables

**Figure S1:** Correlation matrix between performance in each condition (AWD: alternate wetting and drying and CF: continuous flooding) and response variables (response index and slope of the joint regression) for the three traits considered: days to flowering (FL), nitrogen-balance index (NI), and panicle weight (PW). The reference (RP) and progeny (PP) populations are in green and grey, respectively.

**Figure S2**: Single environment and multi-environment (M1 and M2) prediction accuracies in cross validation experiments with 40% of untested entries in the reference population obtained with three statistical models (GBLUP, RKHS-1, RKHS-2). Continuous flooding and alternate wetting and drying water management conditions are in blue and orange, respectively. Three traits are presented: days to flowering (FL), nitrogen balance index (NI) panicle weight (PW). The letters in each panel represent the results of Tukey’s HSD comparison of means and apply to each panel independently. The means differ significantly (p-value < 0 05) if two boxplots have no letter in common.

**Table S1:** Variance components and the associated statistic (F-value for fixed effects and Z-value for random effects) of days to flowering (FL), nitrogen balance index (NI), and panicle weight (PW). Separate analysis of each population and each water management system (alternate wetting and drying - AWD and continuous flooding - CF).

**Table S2:** Variance components and the associated statistic: F-value for fixed effects and Z-value for random effects) of days to flowering (FL), nitrogen balance index (NI), and panicle weight (PW). Separate analysis of each population pooled over water management conditions.

**Table S3:** Variance components for the joint regression for days to flowering (FL), nitrogen balance index (NI), and panicle weight (PW). Results are shown for the reference and progeny populations.

**Table S4:** Mean genomic prediction accuracies in the reference population for the response variables (index and slope) and the performance within each condition (AWD and CF). The results for days to flowering (FL), nitrogen balance index (NI) and panicle weight (PW) are presented. Two statistical models (GBLUP and RKHS) were used.

**Table S5:** Genomic prediction accuracies for across population validation for the response variables (index and slope) and the performance within each condition (AWD and CF). The scenarios used to define the training set are S1 (only the parents), S2 (100 individuals of the RP selected with CDmean) and S3 (the whole RP). Results for days to flowering (FL), nitrogen balance index (NI) and panicles weight (PW) are presented. Two statistical models (GBLUP and RKHS) were used.

**Table S6:** Mean genomic prediction accuracy of the performance within each condition (AWD and CF) using single or multi-environment models in the reference population. For multi-environment models, two methods of cross-validation were used: M1 and M2. Results for days to flowering (FL), nitrogen balance index (NI) and panicle weight (PW) are presented. Two statistical models (GBLUP, RKHS) were used in single environment prediction and three (GBLUP, RKHS-1 and RKHS-2) in multi-environment prediction.

**Table S7:** Genomic prediction accuracies of the performance within each condition (AWD and CF) using single or multi-environment models for across population validation. The scenarios used to define the training set are S1 (only the parents), S2 (100 individuals of the RP selected with CDmean) and S3 (the whole RP). Results for days to flowering (FL), nitrogen balance index (NI) and panicle weight (PW) are presented. Two statistical models (GBLUP, RKHS) were used in single environment prediction and three (GBLUP, RKHS-1 and RKHS-2) in multi-environment prediction.

